# Radiotherapy and cisplatin increase immunotherapy efficacy by enabling local and systemic intratumoral T-cell activity

**DOI:** 10.1101/357533

**Authors:** Paula Kroon, Elselien Frijlink, Victoria Iglesias-Guimarais, Andriy Volkov, Marit M van Buuren, Ton N Schumacher, Marcel Verheij, Jannie Borst, Inge Verbrugge

**Affiliations:** Divisions of Tumor Biology and Immunology, 1066 CX Amsterdam, The Netherlands; Molecular Oncology and Immunology, Oncode Institute, 1066 CX Amsterdam, The Netherlands; Department of Radiation Oncology, The Netherlands Cancer Institute-Antoni van Leeuwenhoek, 1066 CX Amsterdam, The Netherlands

## Abstract

To increase cancer immunotherapy success, PD-1 blockade must be combined with rationally selected treatments. Here, we examined in a poorly immunogenic mouse breast cancer model the potential of antibody-based immunomodulation and conventional anti-cancer treatments to collaborate with anti-PD-1 treatment. One important requirement to improve anti-PD-1-mediated tumor control was to promote tumor-specific cytotoxic T cell (CTL) priming, which was achieved by stimulating the CD137 costimulatory receptor. A second requirement was to overrule PD-1-unrelated mechanisms of CTL suppression in the tumor micro-environment (TME). This was achieved by radiotherapy and cisplatin treatment. In the context of CD137/PD-1-targeting immunotherapy, radiotherapy allowed for tumor elimination by altering the TME, rather than intrinsic CTL functionality. Combining this radioimmunotherapy regimen with low-dose cisplatin improved CTL-dependent regression of a contralateral tumor outside the radiation field. Thus, systemic tumor control may be achieved by combining immunotherapy protocols that promote T cell priming with (chemo)radiation protocols that permit CTL activity in both the irradiated tumor and (occult) metastases.

**Summary statement:** This study reveals that radiotherapy and cisplatin can be ‘re-purposed’ to improve antibody-based immunotherapy success in poorly immunogenic breast cancer by overruling PD-1 unrelated mechanisms of T cell suppression in the tumor micro-environment.

## Introduction

Cancer immunotherapies include adoptive T-cell therapy, therapeutic vaccination, and/or antibody-based immunomodulation. From a technical perspective, antibody-based immunomodulation is relatively straightforward, since immunomodulatory antibodies can essentially be delivered in the same way as conventional anti-cancer drugs. The immunomodulatory antibodies that have been approved for cancer immunotherapy are designed to target the T-cell coinhibitory receptors PD-1 or CTLA-4, and both single or combined treatment induces durable responses in up to 32% of patients with solid tumors (Iwai et al., 2017). Still, the majority of patients do not benefit from this treatment approach (Sharma et al., 2017). Compared to targeting CTLA-4, targeting PD-1 is generally more successful and is associated with fewer autoimmune symptoms (Buchbinder and Desai, 2016). Therefore, targeting PD-1 currently serves as the backbone for developing new combination therapies in order to improve patient response rates. To choose combinations rationally, insight into their combined mechanism of action is required.

Successful immunotherapy relies on the activity of T cells. Upon activation by tumor-specific antigens, CD8^+^ T cells develop into CTLs that can recognize tumor-derived (intracellular) peptides that are presented on the cell surface by MHC class I molecules. As MHC class I molecules are expressed on virtually all body cells, CTLs can in principle target any cancer type. CD4^+^ T cells also promote anti-tumor immunity, either by direct cytotoxic activity or by promoting the activity of CTLs and other immune cells (Melssen and Slingluff, 2017). Several groups postulated that successful immunotherapy relies on a tumor-specific T-cell response that is self-sustained by continuous generation of new effector T cells (T cell priming) and support of their activity (Chen and Mellman, 2013; Spitzer et al., 2017). To enable this cycle, the tumor must essentially act as its own ‘vaccine’ by releasing both recognizable antigens and ‘danger’ signals. Dendritic cells (DCs) can then successfully present these antigens to naïve T cells and provide the appropriate costimulatory and cytokine signals needed to induce T cell clonal expansion and effector differentiation. However, the TME often poses a challenge to T-cell activity. For example, in immunogenic tumors that have given rise to a T-cell response throughout their development, negative feed-back mechanisms come into play that reduce effector T-cell functions. These mechanisms include the activity of regulatory T cells (Tregs) and suppressive activity of myeloid cells, stromal cells, and even the tumor cells themselves (Munn and Bronte, 2016). The PD-1/PD-L1 axis is one example of such a negative feed-back mechanism, in which PD-L1 can be expressed on tumor cells and/or other (immune) cell types present in the tumor, thereby inhibiting T cell function via PD-1 (Hui et al., 2017). Thus, successful immunotherapy requires the elimination of such suppressive mechanisms.

Blocking CTLA-4 enables CD28 costimulation (Krummel and Allison, 1995), which may promote new T cell priming and effector T cell activity. Wei et al. recently showed that blocking CTLA-4 and PD-1 promotes the activity of T cells inside tumors in a complementary fashion (Wei et al., 2017). In addition, data suggests that blocking CTLA-4 promotes T cell priming in cancer patients (Kvistborg et al., 2014). However, combined therapy targeting both CTLA-4 and PD-1 is associated with increased autoimmunity (Sznol et al., 2017) and this combination may therefore rather be avoided when developing new immunotherapy strategies. Targeting CD137 (also known as 4-1BB or TNFRSF9) using agonistic antibodies may provide an alternative. This approach is currently in phase 3 clinical trials (Makkouk et al., 2016) and is being tested in combination with PD-1 blockade in phase 1b clinical trials (Tolcher et al., 2017). CD137 is a costimulatory receptor that belongs to the Tumor Necrosis Factor (TNF) receptor family and its signaling promotes the priming and maintenance of CTL responses by delivering pro-survival and other signals to CD8^+^ T cells (Bartkowiak and Curran, 2015).

Both radiotherapy and chemotherapy can affect the T cell response. These regimens induce tumor cell destruction, which leads to release of antigens and ‘danger’ signals (Galluzzi et al., 2017). In principle, these events may lead to new T-cell priming. However, the likelihood that priming will occur without immunotherapy-based assistance is low, given that radiotherapy almost never gives rise to an ‘abscopal’ effect, i.e. regression of a tumor mass outside the radiation field (Abuodeh et al., 2016). It is speculated from mouse models (van der Sluis et al., 2015) that conventional chemotherapeutic drugs may have important immunomodulatory actions in human, but thus far, there has not been a systemic address of this question in the clinic.

Here, we examined the potential of using radiotherapy and a routinely co-applied conventional chemotherapy (cisplatin) to assist immunotherapy in evoking a systemic, tumor-eradicating T cell response. We provide evidence for immunomodulatory actions of radiotherapy and chemotherapy that make tumors permissive to CTL activity. To our knowledge, this is the first preclinical study that demonstrates the feasiliby and efficacy of combining chemo-, radio- and immunotherapy to improve both local and systemic tumor control. These data argue that conventional anti-cancer regimens can be combined rationally with immunotherapy in order to improve systemic tumor control and increase cure rate and patient outcome.

## Results

### Immunotherapy with CD137 agonism and PD-1 blockade promotes T-cell priming

As a model system, we used mice implanted orthotopically into the fat pad with syngeneic AT-3 breast cancer cells that were treated with immunotherapy and/or radiotherapy after the tumor reached >20mm^2^. Standard immunotherapy consisted of a blocking antibody to PD-1 and an agonistic antibody to CD137 (Figure 1A). In this setting, immunotherapy and radiotherapy as individual treatments merely delayed tumor outgrowth, while combined treatment (i.e. radio-immunotherapy, or RIT) cured the majority of the mice (Figure 1B). We have previously shown that tumor control in this setting relies on CD8+T cells (Verbrugge et al., 2014; Verbrugge et al., 2012).

**Figure 1.**
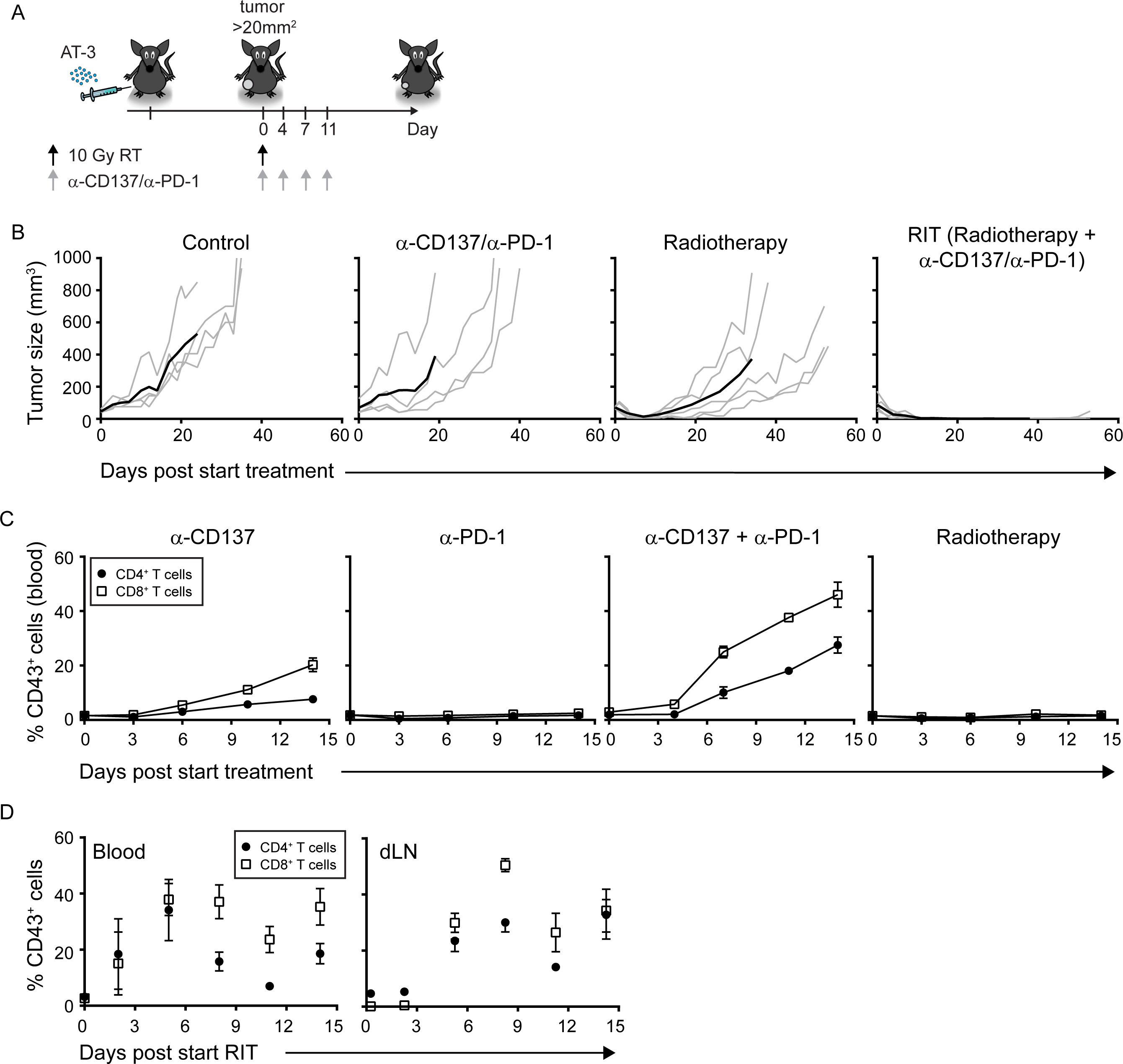
Immunotherapy with CD137 agonism and PD-1 blockade promotes T-cell priming. (A) Experimental set-up. Mice were orthotopically implanted with AT-3 tumor cells and received radiotherapy or immunotherapy alone or in combination after 2–3 weeks when the tumor was established (20 mm^2^). (B) Tumor growth curves measured in mice that received the indicated therapies (n=4–5 mice per group). In each plot, the gray line represents individual mice, and the black line represents the group average. (C) CD43 expression on CD4^+^ and CD8^+^ T cells in blood, as analysed by flow cytometry in tumor-bearing mice (n=5 mice per group) at different time points after the indicated therapy. (D) CD43 expression on CD4^+^ and CD8^+^T cells, measured in the blood (left) and draining lymph node (dLN) of tumor-bearing mice (n=4–5 mice per group), as analyzed at different time points after RIT.

Among single modality treatments, only agonistic antibody to CD137, but not radiotherapy or the blocking antibody to PD-1 induced a T-cell response (Figure 1C), which was measured by the appearance of CD4^+^ and CD8^+^ T cells with a CD43^+^ effector phenotype in blood post treatment (Jones et al., 1994). Adding PD-1 blockade further increased the CD4^+^ and CD8^+^ T-cell responses compared to anti-CD137 agonist treatment alone (Figure 1C). Finally, when immunotherapy with both antibodies was combined with radiotherapy, CD4^+^ and CD8^+^ T-cell responses were also induced, as measured by an increase in effector-phenotype T cells in the blood and in the (inguinal) tumor-draining lymph node (dLN) (Figure 1D). These data suggest that immunotherapy with CD137 agonist antibody promotes T-cell priming, which is increased by PD-1 blockade and is not impeded by concurrent radiotherapy.

### Control of the irradiated tumor by RIT requires T-cell priming

To examine the contribution of newly primed T cells to tumor control, we treated mice with the drug FTY720 that induces the internalization of the sphingosine 1 phosphate receptor 1 (S1PR1). T cells use the S1PR1 to egress from secondary lymphoid organs and the drug prevents them from doing so (Matloubian et al., 2004). RIT was applied while the mice were treated with FTY720 or vehicle (Figure 2A, Figure S2). As a measure of T-cell priming and resulting effector T-cell generation, we measured the percentage of CD4^+^ and CD8^+^ T cells in the dLN that could produce the effector cytokines TNFα and/or IFNγ. RIT increased the percentage of CD8+ effector T cells, and these cells significantly accumulated in the dLN upon FTY720 treatment (Figure 2B, left). In contrast, TNFα-producing CD4^+^ effector T cells were not increased by RIT, nor did these cells accumulate in the dLN upon FTY720 treatment (Figure 2B, right). These data indicate that RIT induced new priming of CD8^+^ T cells and that FTY720 treatment effectively ‘trapped’ these newly primed T cells in the dLN.

**Figure 2:**
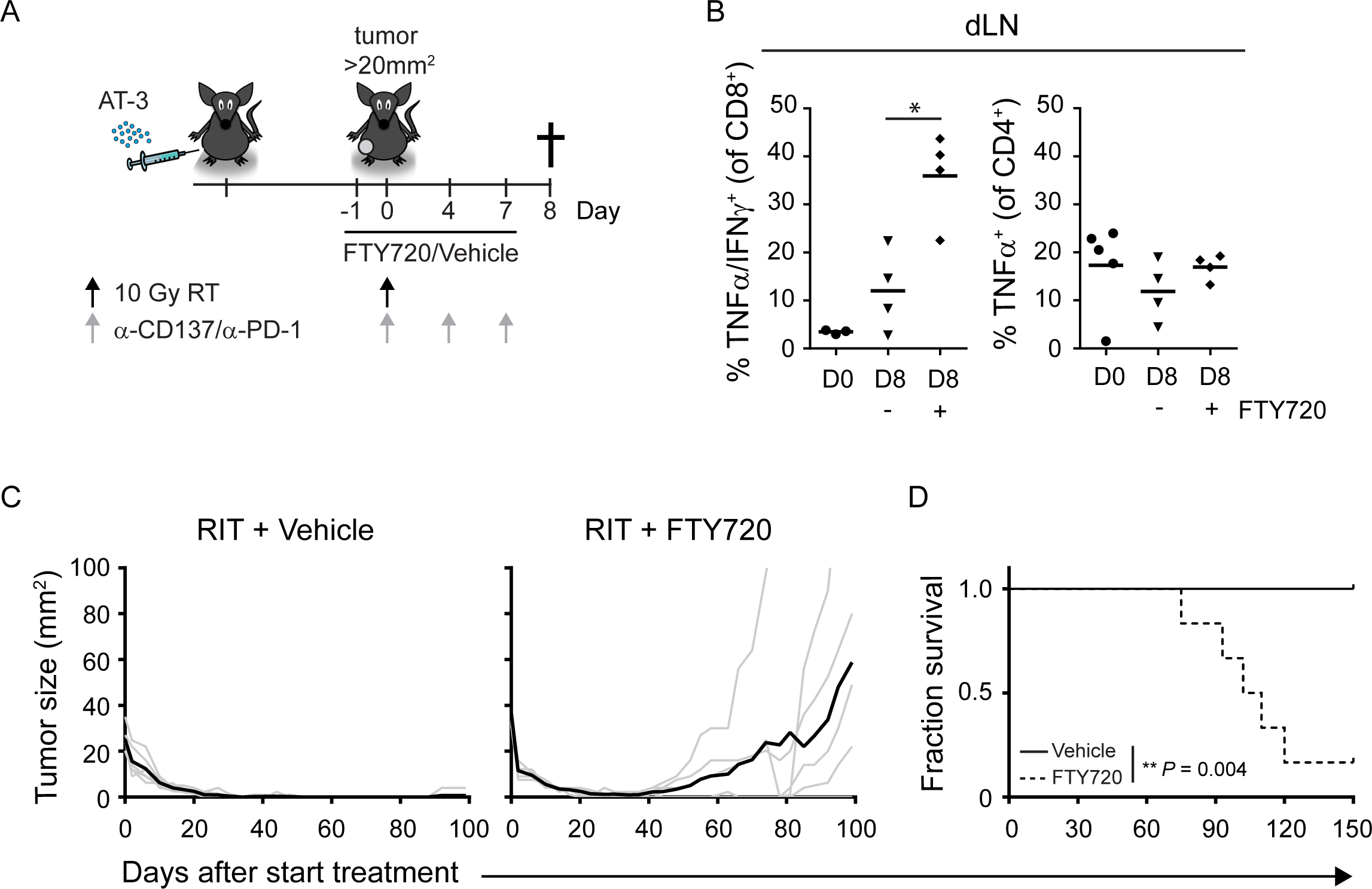
Control of the irradiated tumor by RIT requires T-cell priming. (A) Experimental set-up for B. Mice (n=3–4 per group) were orthotopically implanted with AT-3 tumor cells. After 2–3 weeks, when the tumor was established (20mm^2^), mice received vehicle (saline) or FTY720 in combination with RIT. The horizontal black line indicates the period of FTY720 (or vehicle) administration (3x weekly). On day 8, the animals were sacrificed and analyzed. (B) Summary of the percentage of CD4^+^ (right) and CD8^+^ T cells (left) in the dLN that produce TNFα/IFNy or TNFα in response to *in vitro* re-stimulation, measured before (D0, closed circles) or 8 days after starting RIT (D8). The closed triangles and diamonds at D8 represent mice that received vehicle or FTY720 respectively. (C) Tumor growth curves and (D) survival curves of AT-3 tumor-bearing mice (n=6 per group) receiving RIT as in Figure 1A, where FTY720 (or vehicle) was administered 3x weekly until the end of the experiment.

Importantly, whereas RIT alone led to 100% long-term survival, tumor outgrowth in the RIT-treated mice that received FTY720 was significantly increased (Figure 2C), and overall survival in this group was significantly reduced (Figure 2D). Thus, we conclude that newly primed T cells, raised as a result of RIT make a critical contribution to regression of the irradiated tumor.

### RIT does not induce regression of an abscopal tumor, despite infiltration with newly primed CTLs

Given that RIT induces T-cell priming, we hypothesized that the resulting systemic T-cell response may act against a non-irradiated tumor as well. To examine this, we implanted two separate tumors into the same mouse; one was implanted in the left fat pad and the other was implanted in the contralateral flank. Only the latter tumor was irradiated (Figure 3A). T cell response and tumor regression were examined for both tumors.

We found that the percentage of CD8^+^ T cells among total CD45^+^ (hematopoietic) cells increased significantly in both the irradiated and non-irradiated tumors following RIT (Figure 3B, right). Moreover, the RIT-induced increase of CD8^+^ T cells in the non-irradiated tumor was largely prevented by FTY720 treatment (Figure 3B, right), indicating that this increase was largely due to new T cell priming. The CD4^+^ T cell response following RIT was much less pronounced (Figure 3B). Histological analysis confirmed that CD8^+^ T cells accumulated to a similar extent following RIT in both the irradiated and non-irradiated tumors (Figure 3C). Moreover, infiltration by CTLs - capable of producing effector cytokines IFNγ and TNFα and the cytotoxic effector molecule Granzyme B - was of similar magnitude in the irradiated and non-irradiated tumors (Figure 3D). In contrast, accumulation into tumors of CD4^+^ T cells that could produce TNFα or Granzyme B was not evident (Figure 3E). As compared to immunotherapy alone, RIT increased control of the irradiated tumor. However regression of the non-irradiated tumor did not occur (Figure 3F, Figure S3A) and overall survival of mice (Figure 3G) was not improved. (Hypo)fractionated radiotherapy is more effective than single-dose radiotherapy in enhancing abscopal tumor control by immunotherapy in certain mouse models (e.g. (Dewan et al., 2009; Vanpouille-Box et al., 2017)). However, 3 × 8 Gy (hypo)fractionation did also not enhance immunotherapy-induced control of non-irradiated AT-3 tumors (Figure S3B, C). Thus, CTLs respond to RIT are present in equal measure in the irradiated and non-irradiated tumor, yet only eliminate the irradiated tumor.

**Figure 3:**
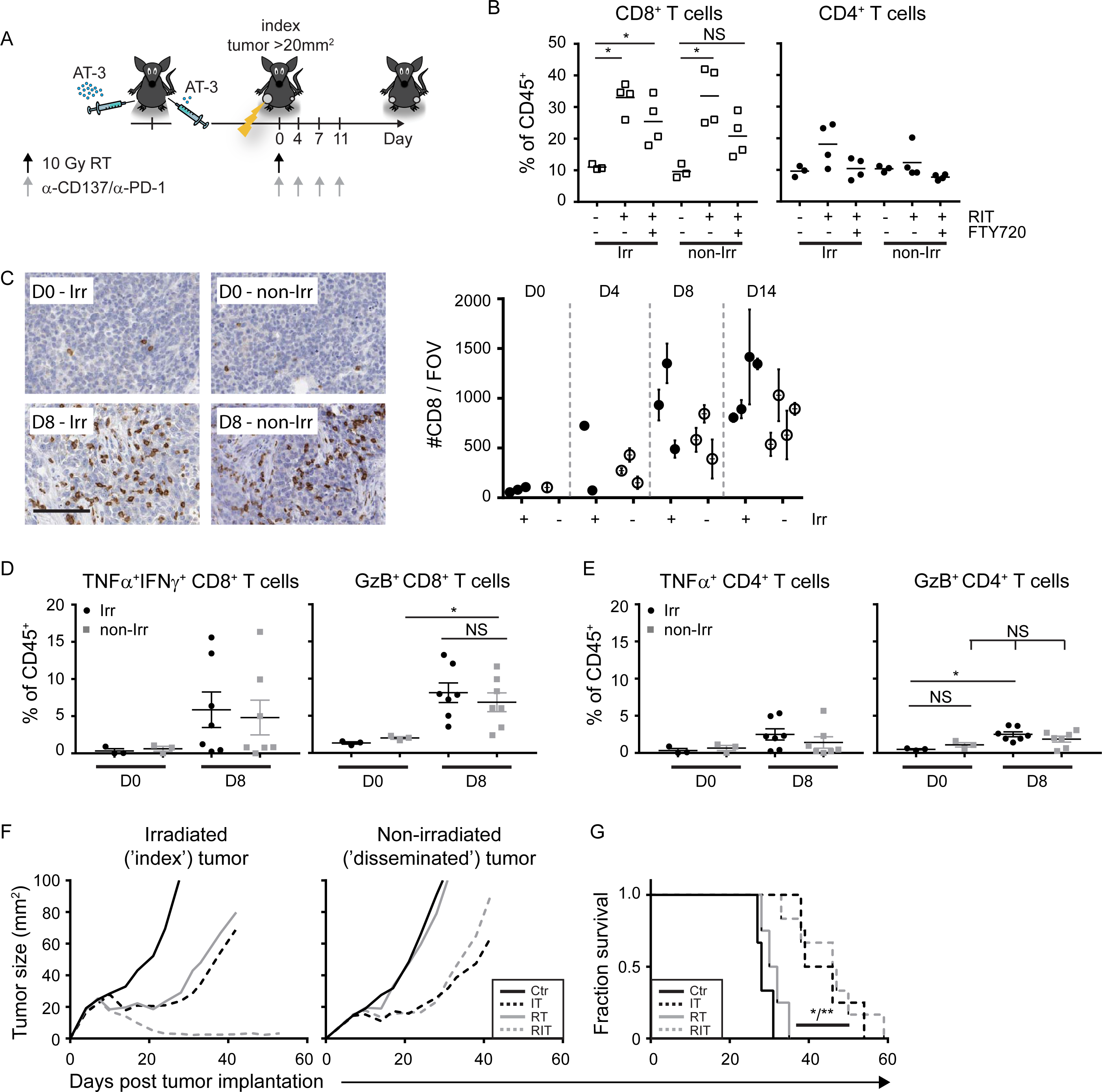
RIT leads to comparable infiltration of irradiated and non-irradiated tumors with T cells, but the non-irradiated tumor does not regress. (A) Experimental set-up. Mice were implanted with AT-3 tumor cells at two sites, orthotopically and in the flank. After 2–3 weeks, when the tumor on the flank was established (20mm^2^), the mice received therapy, wherein only this tumor was irradiated. (B) Summary of the percentage of CD4^+^ and CD8^+^ T cells within the CD45^+^ cell population measured by flow cytometry in the irradiated (Irr) and non-irradiated (non-Irr) tumors before (-) or 8 days after the start of RIT, in the presence or absence of FTY720. (C) AT-3 tumor-bearing mice (n=3 mice per group) were sacrificed before treatment (D0) or 4, 8, or 14 days after the start of RIT and the tumor sections were stained for CD8. Representative images are shown on the left, and the summary data are show on the right. Quantification in the right panel represents the average (± SD) of 5 fields of view (FOVs) for 3 irradiated tumors (filled circles) and 3 non-irradiated tumors (open circles). The images represent 1/6^th^ of a FOV, scale bar = 100 μm. (D-E) Percentage of TNFα^+^IFNγ^+^ or Granzyme B (GzB)^+^ CD8^+^ T cells (D) and CD4^+^ T cells (E) within the CD45^+^ cell population isolated from irradiated and non-irradiated tumors before (D0) and 8 days after the start of treatment. Each symbol represents a single tumor, and the mean is indicated. \**>P*><0.05, Mann-Whitney *>U*> test. (F) Mean tumor growth in mice (n=5–6 mice per group) that receive no therapy (Ctr), radiotherapy (10 Gy, R), either alone or in combination with immunotherapy (IT). (G) Survival curve for mice treated as indicated. \**>P*>< 0.05, between the Ctr and IT groups, \*\**>P*><0.01 between the Ctr and RIT groups and between the R and RIT groups (Mantel-Cox). The difference between the IT and RIT group was not significant (*>P*> = 0.15).

### Tumor-specific T cell priming, neutrophils or macrophages do not limit RIT efficacy

We next addressed a number of potential factors that might disallow RIT-induced CTLs to eliminate the non-irradiated tumor. We first assessed whether the size of the tumor-specific effector CTL pool was a limiting factor. For this purpose, we identified peptide SNPTYSVM from MMTV-Polyoma virus middle-T (PyMT) as an MHC class I-restricted antigen that could raise T cell immunity to AT-3 tumor cells (Figure S4A-F). This enabled us to purposely generate tumor-specific CTL memory *in vivo* by means of vaccinating mice with plasmid (p)DNA encoding this epitope (Figure 4A). Vaccinated mice were challenged as before with two AT-3 tumors and treated with RIT (Figure 4B). Also in this setting, RIT did not enhance control of the non-irradiated tumor (Figure 4C) and did not improve survival of mice (Figure 4D), as compared to immunotherapy alone. These data suggest that the magnitude of the tumor-specific CTL response was not the limiting factor for systemic tumor control following RIT.

**Figure 4:**
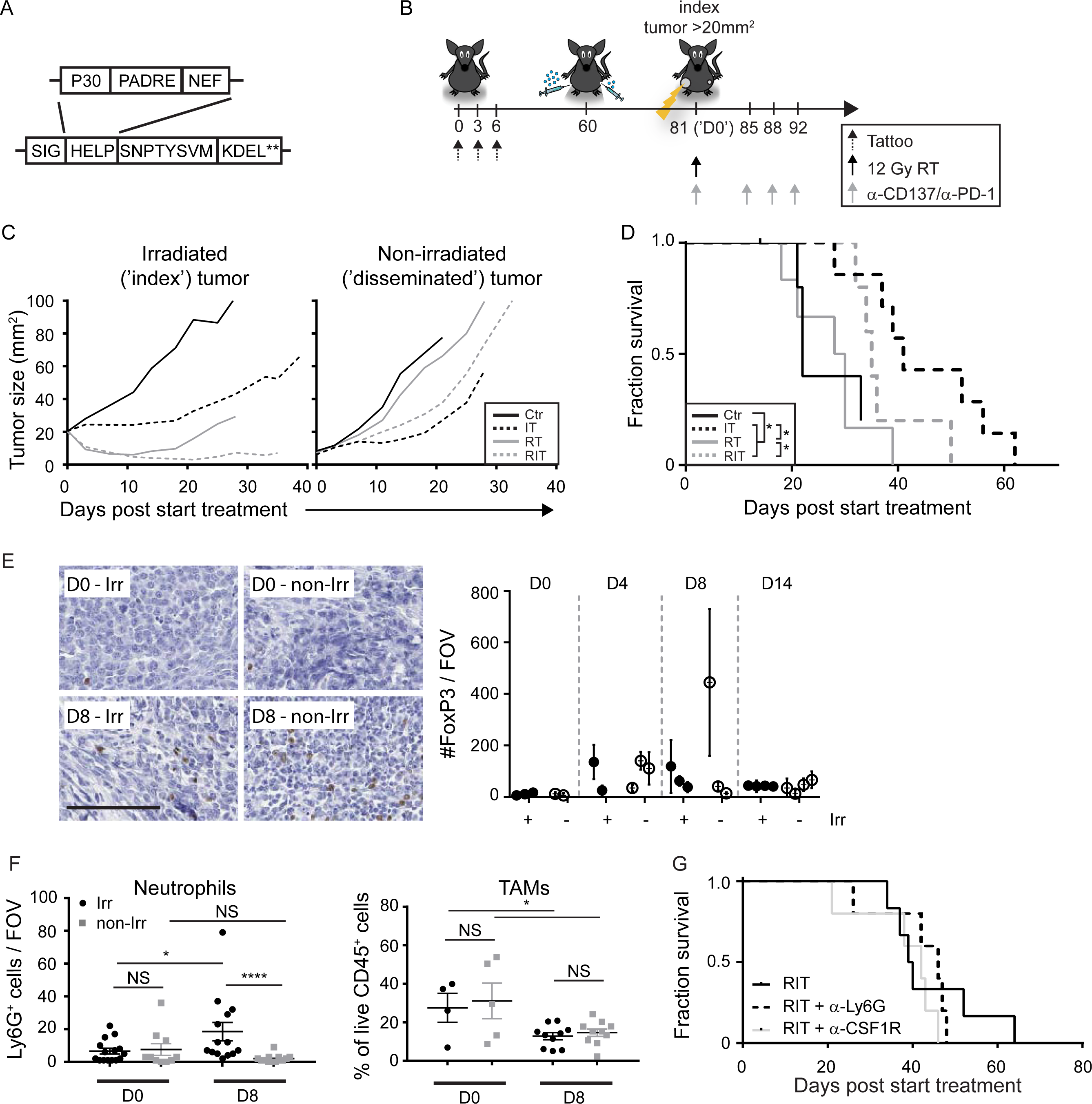
Optimizing tumor-specific T-cell priming, depletion of neutrophils or tumor-associated macrophages is insufficient to control non-irradiated AT-3 tumors following RIT. (A) Vaccine design; the vaccine encodes the H-2K^b^-binding PyMT epitope SNPTYSVM and the human tetanus toxoid P30, PADRE and HIV NEF epitopes. P30 and PADRE bind to mouse MHC II and provide CD4 T cell “help”. **Double stop codon. (B) Experimental set-up. (C and D) Mean tumor size (C) and survival curves (D) of 5–7 mice per group that received the indicated treatments. \**>P*>< 0.05 (Mantel-Cox). (E) FoxP3^+^ regulatory T cells were measured by immunohistochemistry in irradiated (Irr) and non-irradiated (non-Irr) tumors before treatment (D0) and on the indicated days after the start of RIT (n=3 mice per group). Images show representative FoxP3 staining of 10% of the Field Of View (FOV), scale bar = 100 *μ*m. The quantification at the right represents the mean (±SD) of 5 FOVs for 3 irradiated tumors (filled circles) and 3 non-irradiated tumors (open circles). (F) Detection by immunohistochemistry of Ly6G^+^ neutrophils (left) and by flow cytometry of F4/80^+^MHCII^+^ Tumor-Associated Macrophages (TAMs, right) in irradiated (Irr) and contralateral non-irradiated (non-Irr) tumors before (D0) or 8 days after treatment initiation with RIT. Each symbol represents an individual tumor, and the mean ±SEM is shown. (G) Survival curve of tumor-bearing mice that receive RIT in the presence or absence of antibodies targeting Ly6G, or CSF1R (n=5 mice per group).

We next examined which mechanisms of T-cell suppression other than PD-1/PD-L1 interaction may operate in the non-irradiated AT-3 tumors. Treg frequency was low in both the irradiated and non-irradiated tumors and did not change significantly in either tumor following RIT (Figure 4E). Thus, Treg content did not correlate with CTL-mediated tumor control. Tumor-resident neutrophils and macrophages can also locally impair CTL function (Coffelt et al., 2016). RIT increased neutrophil content, as diagnosed by a Ly6G^+^ phenotype, in irradiated tumors, but not in non-irradiated tumors (Figure 4F, left). Thus, neutrophil content positively correlated with CTL-mediated tumor control. RIT decreased macrophage content, as diagnosed by a F4/80^+^MHCII^+^ phenotype (Engelhardt et al., 2012; Franklin et al., 2014) to a similar extent in both the irradiated and non-irradiated tumors (Figure 4F, right). Thus, macrophage content did not correlate with CTL-mediated tumor control. Moreover, effective antibody-mediated depletion of either neutrophils or macrophages (Figure S4G) did not improve control of non-irradiated tumors (Figure S4H), nor did it increase overall survival following RIT (Figure 4G). Taken together, these data suggest that the magnitude of the tumor-specific CTL response, PD-1 signaling, Tregs, neutrophils or TAMs were not key factors that limited CTL activity in the non-irradiated tumor after RIT.

### RIT induces a CTL-permissive TME that is characterized by gene signatures associated with reduced cell proliferation and increased response to tissue injury

RIT led to the same degree of CTL infiltration in the irradiated and non-irradiated tumors, while only the irradiated tumor regressed. This suggests that CTLs can exert their activity on tumor cells only after the tumor has been altered by irradiation. To understand the immunomodulatory effect of irradiation in the context of immunotherapy, we performed mRNA deep sequencing (RNA-seq). Eight days after RIT (to allow sufficient time for T cells to infiltrate both tumors: see Figure 3), we sorted the effector (CD43^+^) CD8^+^ T cells (i.e. ‘CTLs’), CD45^+^ hematopoietic cells (excluding CD43^+^CD8^+^ T cells) and CD45^−^ tumor/stromal cells (Figure 5A). Statistical analysis of normalized read counts revealed the differential expression of 805 genes in CTLs (Figure 5B), 1107 genes in the hematopoietic cells (Figure 5D), and 3045 genes in the tumor/stromal cells (Figure 5F). These genes encode a wide diversity of proteins (Table S1) that perform a multitude of cellular functions.

**Figure 5:**
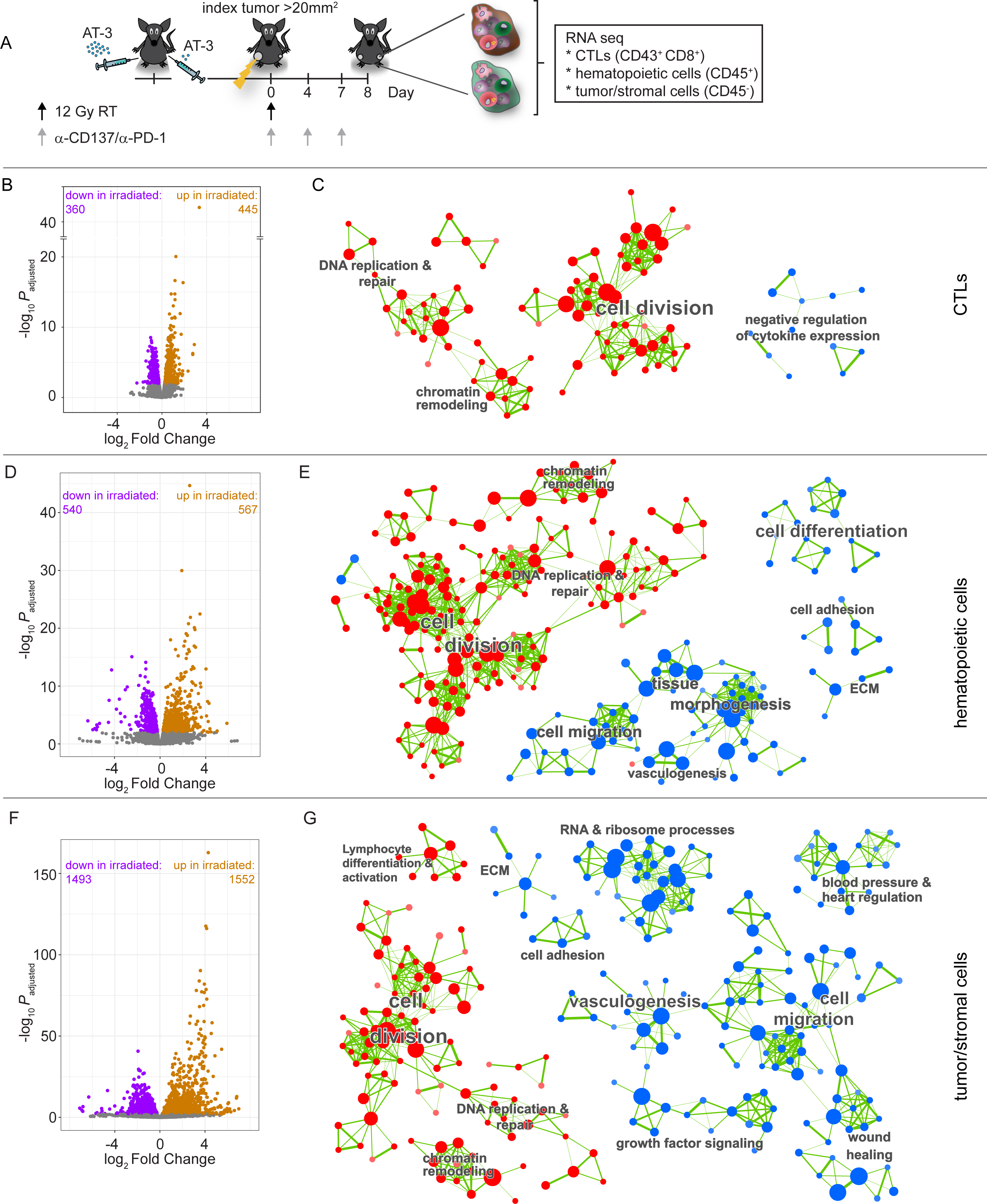
RIT induces a CTL-permissive TME that is characterized by gene signatures associated with reduced cell proliferation and increased response to tissue injury. (A) Experimental set-up: Mice bearing bilateral AT-3 tumors were treated with RIT; on day 8, CD43^+^CD8^+^ CTLs (B, C), CD45^+^ hematopoietic cells (D, E) and CD45^−^ tumor/stromal cells (F, G) were isolated from the irradiated and non-irradiated tumors (n=3 mice per group) and analyzed using RNA-seq. (B, D, and F) Volcano plots of the indicated cell populations showing significant (adjusted P<0.01) transcriptomic changes in the irradiated tumor compared to the non-irradiated tumor. Purple, orange and gray dots represent genes that were downregulated, upregulated or unchanged, respectively. (C, E, and G) Enrichment Maps showing significantly enriched gene sets (biological processes) in blue (enriched in the irradiated tumor relative to the non-irradiated tumor) and red (enriched in the non-irradiated tumor relative to the irradiated tumor). Gene sets that share a high number of genes are clustered together, and the thickness of the green lines represents the number of shared genes. Clusters of similar biological processes are labelled.

Using Gene Set Enrichment Analysis (GSEA) and Enrichment map analysis, we identified groups of biological processes that were differentially modulated between the cell populations at the irradiated and non-irradiated tumor sites. In all three cell populations, gene sets associated with cell division, DNA replication and repair and chromatin remodeling were significantly downregulated in the irradiated tumor (Figure 5C, E, G), congruent with the cells receiving a DNA damaging input in the form of irradiation.

The CTLs in the irradiated tumor included cells that had experienced irradiation, as testified by the differential expression of many genes involved in negative regulation of cell division, DNA replication and repair and chromatin remodeling (Figure 5B,C). The same gene sets were also differentially expressed in other hematopoietic cells (Figure 5D,E) and in non-hematopoietic cells (Figure 5F,G) in the irradiated tumor, as compared to the non-irradiated tumor. In the CTLs, we additionally identified a small group of gene sets associated with negative regulation of cytokine expression (Figure 5C), that included both *Foxp3* and *>IL-10* (Table S1), which may report effects of irradiation. However, we did not identify gene sets associated with increased CTL-intrinsic effector function that could explain the increased CTL efficacy in the irradiated tumor. This finding is consistent with our functional data regarding CD8+ T cells, showing that both irradiated and non-irradiated tumors are infiltrated with effector-phenotype CTLs after RIT (see Figure 3D).

In the hematopoietic and tumor/stromal cells, we identified several biological processes that were significantly different between the irradiated and non-irradiated sites (Figure 5E, G). These included overlapping processes and genes in the hematopoietic and tumor/stromal cells, such as increased cell migration (e.g. *Cxcl17, Cxcl14)*, vasculogenesis (e.g. *Vegfc, Egfl7I)*, and cell adhesion/extracellular matrix (ECM) (e.g. *Selp, Mmp3*, see also Table S1). In addition, and unique to the tumor/stromal cell population, we identified increased expression of gene sets associated with RNA/ribosome processes (e.g.*Rps19, Rps12)* and wound healing (e.g. *Pdgfb, Cxcl12)* in the irradiated tumor as compared to the non-irradiated tumor (Figure 5E, G). In all cell types, we observed increased expression of *Tnf* in the irradiated as compared to the non-irradiated tumor and increased expression of pro-apoptotic *Bax* was observed specifically in the tumor/stromal cells of the irradiated tumor (Table S1).

Taken together, these RNA-seq data clearly reveal the impact of DNA-damaging radiotherapy to the cell populations analyzed by reducing cell division. We did not identify gene sets that were associated with improved CTL-intrinsic functionality in the irradiated tumor. However the TME of the irradiated tumor that permitted CTL activity was clearly different and associated with increased protein translation, blood vessel development and migration of (immune) cells into the damaged tissue, likely reflecting a tissue repair response to irradiation. This gene expression profile in the irradiated TME was associated with increased CTL activity against the AT-3 tumor cells.

### Cisplatin functionally mimics the radiotherapy-induced T cell permissive TME and increases the therapeutic efficacy of RIT

Lastly, we aimed to functionally mimic the generation of the ‘CTL-permissive’ TME induced by radiotherapy in a systemic, body-wide fashion, in order to improve the CTL response to the non-irradiated tumor following RIT. We tested low-dose cisplatin chemotherapy to achieve this effect (Figure 6A) for the following reasons: *i)* Cisplatin has partially the same mode of action as radiotherapy by inducing DNA damage (Dasari and Tchounwou, 2014), *ii)* cisplatin combined with radiotherapy is standard-of-care in the treatment of different types of cancer and *iii)* (low-dose) cisplatin has been shown to support T-cell function (pre-)clinical vaccination studies (van der Sluis et al., 2015).

**Figure 6:**
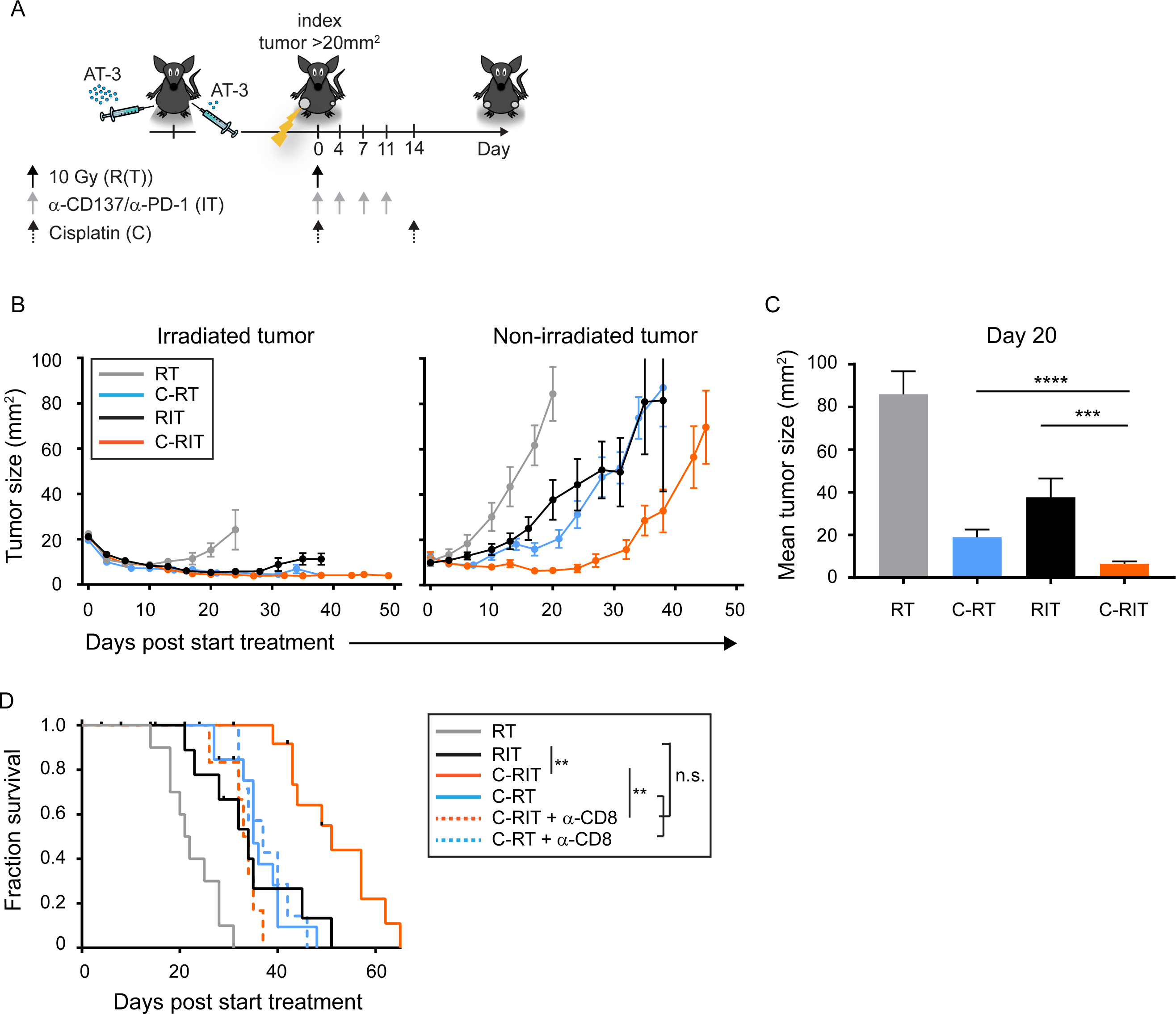
Cisplatin functionally mimics the radiotherapy-induced T cell-permissive TME and increases the therapeutic efficacy of RIT. (A) Experimental set-up: mice bearing bilateral AT-3 tumors were treated with 12 Gy radiotherapy (RT) alone or in combination with anti-CD137/anti-PD-1 mAbs (IT) and/or cisplatin (‘C’; 4 mg/kg). (B) Mean tumor growth (±SEM) of the irradiated and non-irradiated tumors, (C) mean tumor size (± SEM) of non-irradiated tumors on day 20 and (F) Survival curves of the indicated groups of mice; where indicated a CD8-depleting antibody (α-CD8) was administered one day before the start of treatment. Data shown is pooled data from 3 independent experiments of 4–7 mice per group in each experiment. \*\**P* < 0.01; n.s. not significant.

We found that adding cisplatin treatment to RIT improved control of non-irradiated tumors (Figure 6B,C) and increased overall survival (Figure 6D) in a CD8+ T-cell-dependent manner (Figure 6D), despite modestly reducing the magnitude of the T-cell response following RIT (Figure S5). These data indicate that systemic cisplatin treatment functionally mimicked the localized effects of radiotherapy, allowing CTL-mediated growth delay of the non-irradiated tumor and prolonging overall survival following RIT.

## Discussion

There is an unmet clinical need to improve responses to PD-1 blockade, which currently forms the backbone for immunotherapy combinations (Sharma et al., 2017). The PD-1 coinhibitory receptor is associated with tyrosine phosphatase activity that preferentially targets CD28 and thereby inhibits T cell costimulation (Hui et al., 2017). In this way, the PD-1 ‘checkpoint’ can impede both T-cell priming and effector function. However, in cancer patients, PD-1 blockade thus far seems to primarily relieve effector T cells from PD-L1/2-based suppression in the TME (Herbst et al., 2014). Therefore, this approach is likely to be most effective as standalone treatment for immunogenic cancers in which T cells have already infiltrated the tumor (Ayers et al., 2017). Immunotherapy of poorly immunogenic cancers that have not raised a T-cell response will by definition require interventions that induce tumor-specific T-cell priming. However, even in immunogenic cancers that respond to PD-1 blockade alone, new T-cell priming is expected to both strengthen and broaden the anti-tumor immune response, thereby increasing efficacy and combatting resistance (Spitzer et al., 2017). In addition, immune suppression within the TME will pre-exist in immunogenic tumors and may develop in poorly immunogenic tumors once a T-cell response is raised, as a result of negative feedback control on the T-cell response. PD-1/PD-L1 interactions are only a small part of this feedback control, which is exerted by diverse immune- and non-immune cells in the TME. Thus, effective anti-tumor immunity requires both priming of tumor-specific T-cells and a CTL-permissive TME. Here, we show that radiotherapy and conventional chemotherapy can promote intratumoral CTL activity by modulating the TME and by synergizing with an immunotherapy that enables T-cell priming. To our knowledge, this is the first preclinical study that demonstrates the feasiliby and efficacy of combining chemo-, radio- and immunotherapy protocols.

The murine AT-3 breast cancer model we have used carries a foreign antigen, but few T cells are present within AT-3 tumors at steady state (Figure 3C) and PD-1 blockade alone has no therapeutic effect. CD137 agonism induces CTL priming and anti-tumor immunity in this model (Figure 1C). CD137 is expressed on DCs and on activated CD8+ T cells (Figure S1). CD137 triggering on DCs and other myeloid cells can upregulate costimulatory ligands CD80 and CD86 ((Futagawa et al., 2002) and data not shown), which may help to overcome peripheral tolerance and induce T-cell responses to tumor antigens. CD137 triggering on activated CD8+ T cells primary stimulates their proliferation, survival and possibly effector differentiation (Bartkowiak and Curran, 2015), thus supporting CTL priming. In the TME, CD137 agonism can likewise support CTL survival and costimulation by myeloid cells. It can furthermore stimulate hypoxic, CD137-expressing endothelial cells to recruit T cells into the tumor (Palazon et al., 2012). PD-1 blockade incremented CD137-stimulated CTL priming (Figure 1C), indicating that the PD-1 checkpoint can also limit T-cell priming, as observed previously (e.g. (Probst et al., 2005)). We predict that combining PD-1 blockade with any form of immunomodulation that induces CTL priming will be generally very useful clinically. CTLA-4 blockade (e.g. (Vanpouille-Box et al., 2017)) and CD27 agonism (Ahrends et al., 2016; Ahrends et al., 2017) can exert similar effects in distinct tumor models.

Nevertheless, in the combined immunotherapy setting with PD-1 blockade and CD137 agonism, AT-3 tumors were not eliminated, despite a robust CTL response. This problem is also evident in cancer patients that fail to respond to adoptive tumor-specific T-cell therapy (Baruch et al., 2017), highlighting that T-cell suppression in the TME can pose an additional bottleneck for systemic anti-tumor immunity. Importantly, our study demonstrates that radiotherapy can alter the state of the TME to permit effective CTL activity, where PD-1 blockade cannot. Newly primed CTLs raised by our radio-immunotherapy protocol contributed to control of the irradiated tumor. Having a second tumor implanted in the same mouse that was not irradiated allowed us to pinpoint the immune modulating effects of radiotherapy. The non-irradiated tumor was similarly infiltrated by newly primed CTLs, but did not regress, indicating that impediments beyond PD-1 signaling hampered abscopal tumor control. Recent work in a different (CT26) tumor model also revealed that local tumor control by combined radiotherapy and PD-1 blockade was -in part-dependent on newly primed T cells. In that model, control of a simultaneously implanted non-irradiated tumor was also improved by PD-1 blockade (Dovedi et al., 2017). Thus, in that case, PD-1 signaling was the key impediment for CTL activity in the TME, whereas in our AT-3 model, additional impediments were in place. In the AT-3 model, the CTL-enabling effect of radiotherapy could not be reproduced by depletion of neutrophils or TAMs (Figure 4F, G). It has been demonstrated that stimulation of TAMs from PyMT-induced tumors with TLR7/9 agonists (imiquimod, CpG) allows them to reactivate tumor-resident T cells (Engelhardt et al., 2012). TAMs can also phagocytose dead tumor cells and thus enable antigen crosspresentation by DCs. Therefore, altering the functional state of TAMs may be preferred over their depletion to enhance intratumoral CTL activity.

Comparative transcriptome analysis of cell populations from the irradiated and the non-irradiated tumors in the same mice revealed that a ‘CTL-permissive’ TME was associated mostly with changes in CTL-extrinsic, rather than CTL-intrinsic gene signatures. We did not identify gene sets within the CTLs that could explain enhanced efficacy in the irradiated tumor. This indicates that the intrinsic quality of the CTLs that infiltrate the irradiated and non-irradiated tumor after RIT is similar and of good quality, which we also validated by *ex vivo* flow cytometry. Differentially expressed gene sets identified in the CTLs were associated with negative regulation of cytokine production and included *Foxp3* and *Il-10.* We speculate that this is an immune regulatory signature that arose in CTLs that experienced and survived radiotherapy. We cannot exclude that functional activity within this population contributes to enhanced tumor control. Instead, our data indicate that it is mostly CTL-extrinsic parameters that determine CTL efficacy in the irradiated tumor following immunotherapy. Which, and to which extent the differentially expressed gene sets help to permit CTL activity in the irradiated tumor remains speculative. Firstly, increased vasculogenesis function identified in the hematopoietic and tumor/stromal cells within the irradiated tumor could in principle allow more CTLs to infiltrate into the irradiated tumor, which may improve CTL efficacy in tumors where T cell infiltration following immunotherapy is limiting. However, in our tumor model infiltration of CTLs in the non-irradiated and irradiated tumor 8 days after RIT was very similar. Secondly, reduced proliferation of the tumor cells could improve CTL-mediated tumor cell death by allowing T cells more time to complete killing. Thirdly, although we did not identify gene sets associated with increased CTL function, we did identify upregulation of *Tnf* in all cell types within the irradiated tumor. This finding, together with potential sensitization of tumor cells to death receptor induced apoptosis by upregulation of pro-apoptotic *Bax* could contribute to enhanced CTL-mediated tumor control in the irradiated tumor. Interestingly, low-dose cisplatin has also been shown to sensitize tumor cells to CTL-derived TNF-induced cell death (van der Sluis et al., 2015). Fourthly, RNA and ribosome processes were upregulated in tumor/stromal cells of the irradiated tumor. Reits et al. demonstrated that radiotherapy enhances translation in an mTOR-dependent manner, and subsequently increases peptide presentation and tumor cell immunogenicity (Reits et al., 2006). Indeed, mTOR inhibition during RIT treatment reduced therapeutic efficacy (Verbrugge et al., 2014). Finally, processes that are altered non-transcriptionally, which we do not pick with this analysis, may allow increased CTL efficacy in the irradiated tumor.

The use of radio-immunotherapy to treat cancer is a rapidly developing field. Tumor cell destruction by radiotherapy is seen as a mode of vaccination due to the release of antigens and ‘danger’ signals. Thus, the field emphasizes the potential of radiotherapy to contribute to CTL priming, which may result in systemic anti-tumor immunity and ‘abscopal effects’ on non-irradiated tumor masses, when adequately supported by additional interventions (e.g., (Dewan et al., 2009; Rodriguez-Ruiz et al., 2016; Vanpouille-Box et al., 2017; Vanpouille-Box et al., 2015)). Irradiation may help to release danger-associated molecular patterns (DAMPs) such as calreticulin or HMGB1 and/or cytosolic double-stranded DNA that can activate Type I interferon signalling (Vanpouille-Box et al., 2017). Such signals activate DCs changing them from a ‘tolerogenic’ into an ‘immunogenic’ state (Gupta et al., 2012). In tumors that fail to deliver sufficient tumor antigens to DCs *de novo*, radiotherapy-induced debulking of the tumor could furthermore help to reach the ‘antigen threshold’ required for inducing a CTL response. Our study emphasizes that radiotherapy has an additional effect, namely modulating the TME to overcome T cell suppression. To qualify combined effects of immunotherapy and radiotherapy as “abscopal” effects it does not suffice to report an effect on a tumor mass that lies outside of the field of radiation (e.g. (Kwon et al., 2014; Twyman-Saint Victor et al., 2015)), since immunotherapy alone could induce tumor regression. The aim in radio-immunotherapy is to achieve synergy between the two types of treatment. This can only be achieved when T cells that are newly primed as a result of the combined treatment are permitted to exert their activity within the non-irradiated tumor. Here, we show that low-dose cisplatin can facilitate CTL activity in non-irradiated AT-3 tumors in mice treated with PD-1/CD137 targeting therapy, thereby functionally mimicking the immunomodulatory effects of radiotherapy. Based on our findings, ‘re-purposing’ cisplatin at low dose as an immunomodulatory drug, with minimal lymphocyte toxicity may help to convert a CTL suppressive TME into a CTL permissive one, leading to eradication of tumors outside of the field of irradiation. In conclusion, our findings indicate that systemic tumor control may be achieved by combining immunotherapy protocols that promote T cell priming with chemoradiation protocols that permit CTL activity in both the irradiated tumor and (occult) metastases.

## Materials and methods

### Cells

AT-3 cells are derived from the MMTV-Polyoma virus middle-T (PyMT) transgenic mouse, back-crossed to C57BL/6 (Stewart and Abrams, 2007). AT-3 cells were cultured in Dulbecco’s Modified Eagle’s Medium (DMEM), supplemented with 10% fetal calf serum (FCS), 0.1 mM nonessential amino acids, 1 mM sodium pyruvate, 2 mM L-glutamine, 10 mM HEPES, and 30 μM *β* -mercaptoethanol at 37°C, 10% CO_2_.

### Mice

Six-eight-week-old female C57BL/6JRj (B6) mice were obtained from Janvier Laboratories (Le Genest Saint Isle, France) or from in-house breeding within the Netherlands Cancer Institute (NKI) and maintained in individually ventilated cages (Innovive, San Diego, CA) under specific pathogen-free conditions. All mouse experiments were performed in accordance with institutional and national guidelines and were approved by the Committee for Animal Experimentation at the NKI.

### Therapeutic antibodies and reagents

Rat anti-mouse CD137 agonist mAb (clone 3H3, IgG2a) (Shuford et al., 1997) was purified from hybridoma supernatant by affinity chromatography on protein-G. Rat anti-mouse PD-1 mAb (clone RMP1-14, IgG2a) and isotype control (2A3) were purchased from Bio XCell (West Lebanon, NH). FTY720 was purchased from Cayman Chemical (Ann Arbor, MI). Cisplatin (RVG 101430, Pharmachemie BV, Haarlem, the Netherlands) was administered intravenously at 4 mg/kg on day 0 (i.e. immediately after RIT) and on day 14.

### Tumor transplantation and therapy

AT-3 cell transplantation and therapy were performed essentially as described (Verbrugge et al., 2014; Verbrugge et al., 2012), with minor modifications. Briefly, mice were anesthetized with isoflurane, and injected with 1 × 10^6^ AT-3 cells into the fourth mammary fat pad. In some experiments, mice were injected with 0.5 × 10^6^ AT-3 cells into this fat pad on one side and with 2.5 × 10^6^ AT-3 cells on the contralateral flank. The latter tumor was irradiated, and the other tumor served as the non-irradiated ‘abscopal’ tumor. Tumor size was measured using a caliper, and treatment was initiated when the tumors reached 20-25mm^2^. Therapy was done with n=5-10 mice per group. Radiotherapy (RT) was applied using an XRAD225-Cx system (Precision X-Ray, North Branford, CT), as described previously (Kroon et al., 2016; Verbrugge et al., 2014). In brief, the mice were anesthetized with isoflurane and a cone-beam CT scan of the mice was performed. The tumor(s) were localized on the CT scan and targeted with RT at 0.1 mm precision using round collimators 1.0 or 1.5 cm in diameter. A single fraction of 10-12 Gy (225 peak kilovoltage (kVp), filtered with 0.3 mm of copper (3 Gy/min) was delivered. Control mice were anesthetized and underwent a cone-beam CT scan, but were not exposed to RT. Immunomodulatory α-PD-1 and α-CD137 mAbs or an isotype control mAb were diluted in PBS. The antibodies were administered twice weekly for 2 weeks either intraperitoneally (α-PD-1 mAb, 100 μg per injection), or intratumorally (α-CD137 mAb, 25 μg in 10 μl per injection), with the first dose delivered immediately after RT treatment. Tumor transplantation and therapy for RNAseq experiments was performed identically, with the exception that α-CD137 mAb was delivered i.p. (100 μg). The sphingosine-1-phosphate receptor-1 agonist FTY720 was diluted in saline (vehicle) and administered at 2 mg/kg by oral gavage. FTY720 treatment started one day prior to RT and was repeated three times per week throughout the duration of the experiment. All mice were sacrificed when the tumor(s) 2 reached 100-200 mm^2^. A tumor size of 100 mm^2^ was set at a designated end point.

### DNA vaccination

For DNA vaccination, the hair on a hind leg was removed using depilating cream (Veet; Reckitt Benckiser, Slough, UK) on day −1. On days 0, 3 and 6, the mice were anesthetized with isoflurane, and 15 μl of a solution containing 2 mg/ml plasmid DNA in 10 mM Tris and 1mM EDTA, pH 8.0, was applied to the hairless skin with a Permanent Make-up Up Tattoo machine (MT Derm GmbH, Berlin, Germany), using a sterile disposable 9-needle bar with a needle depth of 1 mm and an oscillating frequency of 100 Hz for 45 seconds.

### Flow cytometry

At the indicated time points, tumor-bearing mice were sacrificed and the tumor and tissue were harvested. The tumors were mechanically chopped using a McIlwain tissue chopper (Mickle Laboratory Engineering) and a single-cell suspension was prepared by digesting the tissue in collagenase type A (Roche) and 25 μg/mL DNase (Sigma) in serum-free DMEM medium for 45 min at 37°C. Enzyme activity was neutralized by addition of DMEM containing 8% FCS, and the tissue was dispersed by passing through a 70-μm cell strainer. Single cells were first stained with PE- or APC-conjugated H-2K^b^ PyMT_246-253_ (SNPTYSVM) tetramers for 15 min at 20°C in the dark, followed by fluorochrome conjugated antibodies (see below) for 30 min on ice in the dark in PBS containing 0.5% BSA and 0.01% sodium azide. Intracellular staining following re-stimulation with PMA and ionomycin was performed as described previously (Verbrugge et al., 2014). 7AAD (1:20; eBioscience) or Fixable Viability Dye eFluor^®^ 780 (1:1000; eBioscience) was added to exclude dead cells. All experiments were analyzed using a BD LSRII flow cytometer with Diva software and the generated data were analyzed using FlowJo software.

The following fluorochrome-conjugated mAbs used for flow cytometry were obtained from BD Pharmingen (Breda, the Netherlands) unless otherwise specified: α-CD8-FITC (1:100, clone 56-6.7), α-CD4-eFluor450 (1:200, clone GK1.5), α-TCR*β*-PECy5 (1:200; clone H57-597), α-CD43-PerCPCy5.5 (1:200, clone 1B11 (BioLegend, San Diego, CA)), α-CD45.2-eFluor605 (1:50; clone 30-F11), α-CD4-FITC (1:100, clone GK1.5), α-CD8-V450 (1:300, clone 56-6.7) α-CD11b-AF700 (1:200, clone M1/70), α-CD8-AF700 (1:200, clone 56-6.7), α-IFNy-APC (1:100, clone XMG1.2), and α-TNFα-PECy7 (1:200, clon8e MP6-XT22).

### Prediction of PyMT peptides and generation of PyMT-H-2K/D^b^ multimers

To identify AT-3 tumor antigens, we first used epitope prediction tools to define PyMT-derived peptides that could potentially bind to H-2K^b^ and/or H-2D^b^MHC class I molecules. These peptides were then synthesized by the peptide facility at the NKI, and MHC tetramers were produced by UV-induced peptide exchange as described previously (Toebes et al., 2006). In brief, 28 peptides of PyMT (protein ID: NP_041265.1) predicted to bind either H-2K^b^ or H-2D^b^ (NetPan MHC 3.0 and NetPan MHC 4.0) were synthesized by the peptide facility at the NKI. These peptides were individually exchanged into H-2K^b^ or H-2D^b^ molecules that had been refolded with a UV-sensitive peptide, allowing the generation of monomers with multiple specificities via a single reaction (Toebes et al., 2006). The resulting monomers were subsequently multimerized and conjugated to phycoerythrin (PE) or allophycocyanin (APC) and then used to screen for T cell reactivity to MHC I restricted PyMT epitopes using flow cytometry.

### RNA preparation and sequencing

Using flow cytometry, CD43^+^ CD8^+^ T cells (‘CTLs’), CD45+hematopoietic cells, and CD45^−^ (tumor/stromal) cells were isolated from both the irradiated and non-irradiated tumors of 9 mice per experimental group, and material from 3 mice was pooled per sample to retrieve sufficient RNA. Cells were collected in RLT lysis buffer (QIAGEN) and total RNA was extracted using the RNAeasy mini kit (QIAGEN) in accordance with the manufacturer’s instructions. The integrity of the total RNA was assessed using a 2100 Bioanalyzer System (Agilent). Only RNA samples with an RNA Integrity Number (RIN) > 8 were used to create the library. Poly-A-selected RNA libraries were prepared using the TruSeq RNA library protocol (Illumina) and the resulting libraries were sequenced using an Illumina HiSeq2500 with V4 chemistry, with 50-bp single-end reads per lane.

### Transcriptomics analysis of illumina sequencing data

Sequencing reads in FASTQ files were mapped to the mouse genome (build GRCm38.77) using Tophat v2.1 (Trapnell et al., 2009), and the read summarization program HTseq-count (Anders et al., 2015) was used to count uniquely mapped reads against annotated genes. Differential expression analysis was performed using the DESeq package in R (Love et al., 2014). P-values were corrected for multiple comparisons, based on the False Discovery Rate (FDR), with significance considered at a *q*-value <0.01. Volcano plots were generated using ggplot2 (https://www.springer.com/gp/book/9780387981413).

Normalized read counts were used as input for Gene Ontology Gene Set Enrichment Analysis (GSEA) version 3.0 (Mootha et al., 2003; Subramanian et al., 2005) to identify groups of biological processes that were differentially expressed between cell populations obtained from the irradiated site and cell populations obtained from the non-irradiated site. We used the MSigDB C5 collection to identify enriched gene ontology (GO) biological processes (BP). GSEA was performed with default parameters and gene set permutations. To gain a better overview of the linked biological processes, we generated enrichment maps using the Enrichment Map app v3.1.0 (Merico et al., 2010), using cut-off values set at Q = 0.1 and Jaccard Overlap Combined = 0.375. We illustrated the biggest gene set clusters and manually assigned the more general processes that these clusters represent.

The RNA-seq data reported in this paper have been deposited in the ArrayExpress database at EMBL-EBI (www.ebi.ac.uk/arrayexpress) under accession number E-MTAB-6914.

### Immunohistochemical analysis

Harvested tumors and tissues were fixed for 24 h in ethanol (50%), acetic acid (5%), and formalin (3.7%), embedded in paraffin, and then sectioned randomly at 5 μm. The sections were then stained as described previously (Kroon et al., 2016). In brief, fixed sections were rehydrated and then incubated with primary antibodies to CD8 (eBioscience; clone 4SM15) and Foxp3 (eBioscience; clone FJK-16s). Endogenous peroxidases were blocked with 3% H_2_O_2_ and the sections were then incubated with biotin-conjugated secondary antibodies, followed by incubation with HRP-conjugated streptavidin-biotin (DAKO). The substrate was developed using diaminobenzidine (DAB) (DAKO). We included negative controls to determine background staining, which was negligible. The stained sections were digitally processed using an Aperio ScanScope (Aperio, Vista, CA) equipped with a 20x objective. ImageJ software was used to quantify the number of positive cells in 3-5 random fields of view (FOV) per slide.

### Statistical Analysis

All summary data were analyzed using GraphPad Prism version 6 (GraphPad Software, La Jolla, CA). Differences between various treatment groups were analyzed using the Mann-Whitney *U* Test. Differences in survival curves were analyzed using the Log Rank (Mantel-Cox) test. Differences with P-value <0.05 were considered statistically significant.

## Author contributions

PK, VI, EF, and IV designed and performed the *in vitro* and *in vivo* experiments. EF and AV analyzed the RNA sequencing data. MvB and TNS contributed scientifically and technically to the identification of an AT-3 tumor antigen and the generation of MHC multimers. IV acquired funding and conceptualized the study with MV and JB, and co-supervised the study with JB. IV and JB wrote the paper with help of PK, EF, VI and AV. All authors agreed to the final version of the manuscript.

## Acknowledgments

We thank the personnel of the animal facilities for mouse husbandry and Jan-Jakob Sonke and Artem Khmelinskii for their expertise in the small animal radiotherapy facility at the Netherlands Cancer Institute. We thank the Intervention Unit at the Netherlands Cancer Institute for help with the mouse experiments. We thank Ron Kerkhoven, Arno Velds, Marja Nieuwland, and Iris de Rink of the NKI genomics core facility for providing advice, technical assistance and bioinformatics support regarding analyzing the RNA sequencing experiments. We thank Hideo Yagita for providing the anti-CD137 (3H3) antibody. We thank Mireille Toebes for providing expert advice and technical assistance in generating tetramers. We thank Henk Hilkman and Dris El Atmioui at the Peptide Production Facility for generating the peptides. We thank Julia Walker and Nelleke Goense for providing technical assistance. We also thank Karin de Visser and all members of the Borst laboratory for helpful discussions. We thank Maries van den Broek for critical reading of the manuscript. This work was financially supported by the Dutch Cancer Society to IV (grants NKI 2013-5951 and 10764) and a Health Holland public-private partnership grant in collaboration with Elekta to Jan-Jakob Sonke (grant LSHM15036). Jannie Borst is listed as an inventor on a patent on CD27-agonist antibodies and is in receipt of research grants from Aduro Biotech Europe, which is unrelated to the work described in this manuscript. The authors have no additional financial interests.

## Non-standard abbreviations

DC: Dendritic cell
dLN: draining lymph node
Gy: Gray
PyMT: Polyoma virus Middle T
RNAseq: RNA sequencing
R(I)T: Radio-(immuno)therapy
S1PR1: Sphingosine 1 phosphate receptor 1
TAM: Tumor-associated macrophage
TME: Tumor micro-environment
Treg: Regulatory T cell

